# Attentional modulation of orthographic neighborhood effects during reading: Evidence from event-related brain potentials in a psychological refractory period paradigm

**DOI:** 10.1101/337238

**Authors:** Milena Rabovsky, Markus Conrad, Carlos J. Álvarez, Jörg Paschke-Goldt, Werner Sommer

## Abstract

It is often assumed that word reading proceeds automatically. Here, we tested this assumption by recording event-related potentials during a psychological refractory period (PRP) paradigm, requiring lexical decisions about written words. Specifically, we selected words differing in their orthographic neighborhood size – the number of words that can be obtained from a target by exchanging a single letter – and investigated how influences of this variable depend on the availability of central attention. As expected, when attentional resources for lexical decisions were unconstrained, words with many orthographic neighbors elicited larger N400 amplitudes than those with few neighbors. However, under conditions of high temporal overlap with a high priority primary task, the N400 effect disappeared. This finding indicates strong attentional influences on the incidental processing of orthographic neighbors during word reading, providing novel evidence against the automaticity of processes involved in word reading. Furthermore, in conjunction with the observation of an underadditive interaction between stimulus onset asynchrony (SOA) and orthographic neighborhood size in lexical decision performance, commonly taken to indicate automaticity, our results raise issues concerning the standard logic of cognitive slack in the PRP paradigm.

## 1 Introduction

Word reading is a highly overlearned every-day activity, and is widely assumed to proceed automatically (see e.g., the vast majority of textbooks). Empirically, the automaticity of word processing receives support from phenomena like the Stroop effect (Stroop, 1935), where the meaning of color words is accessed even when this interferes with the task of naming the color in which the word is printed (see also Augustinova & Ferrand, 2014). Furthermore, automatic word processing is also implicitly assumed in many models of visual word recognition (e.g. Coltheart et al., 2001; Forster, 1976; McClelland & Rumelhart, 1981; Jacobs & Grainger, 1996; Morton, 1969; Plaut et al., 1996; Seidenberg & McClelland, 1989). However, recent evidence from the *psychological refractory period (PRP)* paradigm, which is well-suited to investigate attentional influences on cognition (e.g. Keele, 1973; Pashler, 1984; Pashler & Johnston, 1989; Schweickert, 1978; Telford, 1931; Welford, 1952) has casted doubt on this assumption, triggering considerable debate about the automaticity of visual word processing (Allen et al., 2002; Besner, Risko, Stolz, White, Reynolds, O’Malley, & Robidoux, 2016; Cleland et al., 2006; Lien et al., 2006; Lien et al., 2008, McCann et al., 2000; O’Malley et al., 2008; Reynolds & Besner, 2006; Rabovsky et al., 2008; Ruthruff et al., 2008; Vachon & Jolicoeur, 2012). The present experiment combined a PRP paradigm with recordings of event-related potentials (ERPs) to further elucidate this question. In particular, we focused on attentional modulations of one sub-process occurring during lexical processing, namely the incidental processing of orthographic neighbors (ON) during word reading. In the following, we will first provide background on how the PRP paradigm can be combined with ERPs and review how attentional modulations of word processing have been investigated in PRP studies. Finally we will introduce the focus of the present study, orthographic neighborhood size.

### 1.1 Combining PRP paradigm and ERPs

In a typical PRP experiment, two stimuli requiring separate responses are presented in rapid succession. The delay between the stimuli, known as *stimulus onset asynchrony (SOA)*, is varied across trials. Whereas response times to the first stimulus (RT1) are relatively unaffected by SOA, response times to the second stimulus (RT2) are longest at short SOA and decrease with increasing SOA to a point after which RT2 remains constant. The standard account for this pattern of results is to assume that some cognitive processes need a mechanism which can only be dedicated to one task at a time, producing a processing *bottleneck*. Although models assuming a bottleneck differ in their assumptions about the processing stage at which it is localized, there is considerable evidence in favor of a central bottleneck involving decision and response selection processes (McCann & Johnston, 1992; Pashler, 1984; Pashler, 1989; Pashler & Johnston, 1989). According to these models the first task (T1) enters the bottleneck first and is therefore unaffected by the degree of overlap with the second task (T2). However, processes of T2 that need the bottleneck mechanism are postponed while it is occupied by T1. The resulting waiting period for T2, termed *cognitive slack*, increases with decreasing SOA; this explains the effect of SOA on RT2. Processes before and after the bottleneck stage(s) are assumed to operate in parallel for both tasks without interference.

An important and non-trivial prediction of bottleneck models is that increasing difficulty – and thus time demands – of T2-related processes that are functionally localized prior to the bottleneck should have less impact on RT2 when the SOA is short than when it is long; in case of short SOAs T2 slowing takes place during the waiting period and hence is *absorbed into slack* (see Fig. 1, bottom). The effects of difficulty manipulations of T2-related pre-bottleneck processes should therefore decrease with decreasing SOA; in other words, processing delays due to task overlap (SOA) and difficulty of T2 pre-bottleneck processes should combine *underadditively* (Pashler, 1984; Pashler & Johnston, 1989; Schweickert, 1978). In contrast, the slowing induced by increased difficulty of T2 related processes that take place during or after the central bottleneck will result in RT2 slowing that is independent of SOA, because the slowing cannot be absorbed into the waiting period (see Fig. 1, top). Thus, the effects of processing difficulty of T2-related bottleneck processes and task overlap on RT2 combine *additively* (Pashler, 1984; Pashler & Johnston, 1989; Schweickert, 1978). Based on these predictions, following the logic of cognitive slack (McCann & Johnston, 1992; Schweickert, 1978), it is possible to investigate whether a mental process of interest depends on the availability of the central bottleneck, if this process is included in T2 of a PRP experiment, and its difficulty is manipulated.

**Figure 1.**
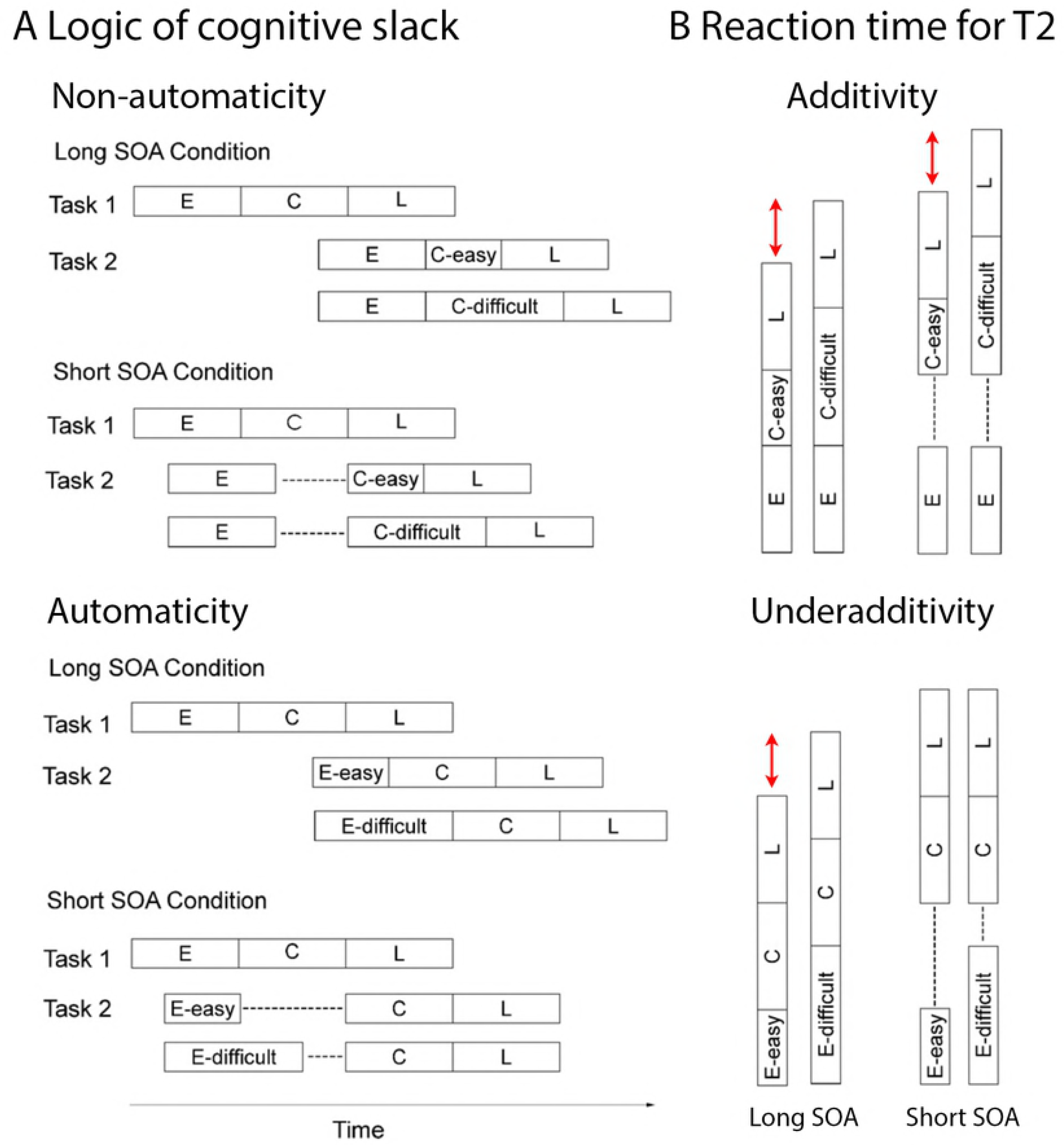
A. Schematic depiction of the logic of cognitive slack. Early (E), central (C), and late (L) processes of Task 1 (T1) and Task 2 (T2) of a PRP paradigm, for long and short stimulus onset asynchrony (SOA) conditions. Top. Difficulty manipulation of a central T2 process, which is non-automatic, i.e. which needs the central bottleneck to proceed, with the difficulty effect predicted not to differ between SOA conditions. Bottom. Difficulty manipulation of an early T2 process, which is automatic, i.e. located before the central bottleneck, with the difficulty effect predicted to be absorbed into slack (- - -) in the short SOA condition. B. Predicted reaction times (RT) for T2 (extracted from panel A and synchronized to the start of T2), for the long SOA condition on the left and the short SOA condition on the right. The key prediction is highlighted in red. Specifically, the difficulty manipulation of a non-automatic process (top) is predicted to have the same effect on RT in the short and long SOA condition. On the other hand, the difficulty manipulation of an automatic process (bottom) is predicted to have an effect in the long SOA condition, but this effect should be eliminated (or diminished) in the short SOA condition.

Johnston et al. (1995) labeled the central bottleneck mechanism, which can only be dedicated to one task at a time, *central attention*, distinguishing it from *input attention*, which is assumed to limit more peripheral forms of processing. Both forms of attentional limitations appear to be functionally independent (McCann & Johnston, 1992). Following Johnston et al. (1995), the SOA manipulation in the PRP paradigm has often been understood as a manipulation of the availability of central attention for T2 processing. Therefore, additive or underadditive combinations of the effects of SOA and the difficulty manipulation of a given process of T2 have been interpreted to indicate whether or not that process depends on the availability of central attention. It is important to note that other accounts of central attention, which characterize it as a scarce resource that can be gradually distributed across tasks (Kahnemann, 1973; McLeod, 1977; Tombu & Jolicoeur, 2003), can predict the same pattern of results. If T1 is performed with priority, as required in most PRP experiments, one might assume that participants primarily devote all their attentional resources to this task, although in principle a more graded resource sharing might be possible as well (Tombu & Jolicoeur, 2003).

Combining the PRP paradigm with ERP recordings can provide important additional insights into attentional modulations of cognitive processes because ERPs provide a continuous on-line measure of the processes between stimulus presentation and response and are therefore well-suited for investigating the processes taking place during cognitive slack (see section ‘Word processing in the PRP paradigm’, below, for details). In the PRP paradigm, ERPs elicited by the T2 stimuli overlap with the ERPs to stimuli and responses of T1. But, as electric fields of several sources combine linearly without interacting (e.g. Nuñez, 1981), it is possible to isolate the effects of a particular experimental factor by a subtraction procedure, which eliminates all invariant overlapping activity. In our study, for example, the effect of ON on the ERP was isolated by subtracting the ERPs to words with many ON from those to words with few ON within each experimental condition. This way, it was possible to identify for each SOA the effect of ON in terms of ERP polarity, latency, and amplitude. Pioneered by Osman and Moore (1993), several previous studies have used this approach, and more recently also applied it to language processing (e.g., Hohlfeld, Mierke, & Sommer, 2004; Hohlfeld, Sangals, & Sommer, 2004; Lien et al., 2008; Martin-Loeches et al., 2009; Rabovsky et al., 2008; Sommer & Hohlfeld, 2008).

### 1.2 Word processing in the PRP paradigm

Several studies have investigated the dependency of visual word processing on central attention with the PRP paradigm, most often by using a visual lexical decision task (LDT) as T2. In a visual LDT, participants decide whether a presented letter string is a real word or not by pressing one of two buttons; this decision requires lexical processing. In PRP studies, the difficulty of lexical processing has often been manipulated by means of word frequency. It is well-established that high frequency words are processed faster and more accurately than low frequency words (Forster & Chambers, 1973; Howes & Solomon, 1951; Rubenstein et al., 1970). This finding has often been taken to indicate that word frequency influences lexical processing in visual word recognition (Broadbent, 1967; Forster & Chambers, 1973; Monsell, 1991; Monsell et al., 1989; Morton, 1969), with lexical processing being more difficult for low frequency words. If lexical processing is automatic – that is, independent of central attention – the effect of word frequency should interact underadditively with SOA. Results have been somewhat mixed, but the overall picture emerging from a number of behavioral studies shows predominantly underadditive interactions between word frequency and SOA (more pronounced in older as compared to younger adults; Allen et al, 2002; Lien et al., 2006) if experimental power is sufficient (Cleland et al., 2006; McCann et al., 2000). These results suggest that lexical processing does not require central attention. However, please note that complete automaticity should not just result in underadditivity, but in a complete disappearance of the effects at short SOA if the slack period is long enough, which is the case in most PRP experiments (please see Fig. 1). Underadditive interactions can also occur when the effects at short SOA are still present but smaller than at long SOA; this is the pattern in the PRP studies discussed above, and can be seen as suggesting partial automaticity.

Importantly, underadditive interactions between SOA and experimental manipulations affecting a cognitive process of interest cannot be unambiguously interpreted based on behavioral data alone and the continuous online signal provided by ERPs can provide important disambiguating information. In the context of word processing, the N400 component in the ERP is of special interest. The N400 is a negative deflection at centroparietal electrode sites peaking around 400 ms after a potentially meaningful stimulus (Kutas & Federmeier, 2011). While the specific functional basis of N400 amplitudes is still controversial it seems clear that the N400 reflects aspects of (lexico-)semantic processing (for a review of verbally descriptive theories see Kutas & Federmeier, 2011; for computational accounts see Frank, Otten, Galli, & Vigliocco, 2015; Laszlo & Armstrong, 2014; Laszlo & Plaut, 2012; Rabovsky, Hansen, & McClelland, 2016; Rabovsky & McRae, 2014). Both word frequency and lexicality (i.e., the difference between words and pseudowords) affect N400 amplitudes (e.g., Braun et al., 2006; Barber et al., 2004) and both effects have been interpreted as electrophysiological indicators of lexical processing.

Capitalizing on these electrophysiological indicators to examine attentional modulations of lexical processing in more detail, Rabovsky et al. (2008) recorded ERPs in a PRP study with a visual LDT as T2 and the difficulty of lexical access being manipulated by means of word frequency. The set-up for this study was basically the same as for the current experiment, with the exception that the variable of interest was changed from word frequency in the previous study to orthographic neighborhood in the current experiment. Replicating earlier findings (e.g., Cleland et al., 2006), they found an underadditive interaction of word frequency and SOA in performance, suggesting that lexical processing could proceed while central attention was dedicated to another task. However, if lexical processing was completely automatic, ERP effects of word frequency and lexicality should have been completely independent of task overlap, that is, SOA, and this was not the case. Instead, both effects were moderately delayed and reduced in amplitude in the short as compared to the long SOA condition (see also Lien et al., 2008). The delay of the ERP effects was much less pronounced than the delay of RT2, suggesting that lexical processing was not halted while the central bottleneck was occupied by T1 (as was the case for decision and response selection processes involved in the LDT). Instead, the results were taken to suggest that, while not completely precluded by unavailable central attention, lexical processing was slowed and associated with less cognitive activity in conditions of high task overlap (for similar interpretations concerning slight delays of ERP effects in non-linguistic PRP studies see Luck, 1998; Arnell et al., 2004; Dell’Aqua et al., 2005; Brisson & Joliecoeur, 2007). Following De Jong (1995), Rabovsky et al. (2008) suggested that this modification of lexical processing under conditions of high task overlap could be due to participants inhibiting visual word processing structures via top-down control in order to optimize T1 performance. This would result in less responsive processing units and less efficient processing in the visual word processing system.

### 1.3 The present study

The present study aimed to further explore attentional modulations of lexical processing, combining ERP recordings with the PRP paradigm. Here, we focused on a particular sub-process involved in visual lexical processing namely the incidental processing of a word’s orthographic neighborhood (ON). ON size is defined as the number of words that can be obtained from a target by exchanging a single letter, preserving letter positions (Coltheart et al., 1977). Typically, large ONs facilitate ‘word’ responses and impair ‘non-word’ responses in lexical decision tasks (Andrews, 1989; Forster & Shen, 1996; Grainger & Jacobs, 1996; Sears et al., 1995). These results have been explained by assuming that lexical decisions are (at least partly) based on the summed activation across all word representations; the summed activation should be higher if a word activates many ONs, facilitating the acceptance of words but impairing the rejection of pseudowords (Grainger & Jacobs, 1996).

In addition, large ONs have been found to reliably enhance the amplitude of the N400 component to both words and pseudowords (Holcomb et al., 2002; Laszlo & Federmeier, 2011). ON effects on ERPs are assumed to reflect increased activation in the lexical semantic system induced by stimuli with large ONs, due to the activation of lexical (e.g. Braun et al., 2006) or semantic (Holcomb et al., 2002; Laszlo & Federmeier, 2011; Laszlo & Plaut, 2012; Müller et al., 2010) representations not only of the presented word but also of its ONs. Alternatively, some researchers have suggested that the large N400 for stimuli with many ON might reflect inhibition of co-activated representations (Debruille, 2007; Molinaro et al., 2010; see Discussion section for more details). In short, ON effects on performance and ERPs obtained in single task contexts are often seen as indicators of the amount of activation (and/ or inhibition) in the lexical semantic system, making them an interesting tool to investigate how lexical semantic processing is modulated by the availability of central attention.

In a behavioral PRP study, Reynolds and Besner (2006) investigated ON effects on reading aloud, which have also been found to be facilitating in single task contexts (Carreiras et al., 1997; Sears et al., 1995) and reported additive effects of SOA and ON. ON effects during reading aloud were taken to reflect lexical contributions to phonological recoding; based on their results the authors suggested that lexical contributions to phonological recoding during reading aloud cannot take place when attention is focused on another task. Since ON effects are strongly modulated by task context (see Discussion section) it is of interest whether the processes underlying ON effects are similarly affected by attention during lexical decisions.

Aiming to investigate the (non-)automaticity of ON-related lexical processing during visual lexical decisions, the present study combined ERP recordings with a PRP paradigm that involved a high-priority pitch discrimination T1 and a visual lexical decision T2 about written words with either many or few ON. Following standard logic of cognitive slack (see Fig. 1 and section 1.1, above), an underadditive interaction between behavioral influences of ON and SOA indicates automaticity of ON-related lexical processing while an additive combination of the effects of SOA and ON indicates that the respective processes require central attention. Influences of task overlap on enhanced N400 amplitudes for words with many ON, which provide a continuous online indicator of ON-related activity were examined in order to more fully understand how lexical processing is modulated by the availability of central attention. Specifically, if ON-related lexical processing is completely automatic, ON effects on N400 amplitudes should not differ between the short and long SOA condition. On the other hand, amplitude reductions or delays of the N400 effects indicate non-automaticity.

## 2 Results

### 2.1 Performance

#### Task 2

Mean correct RTs for lexical decisions are displayed in Figure 2. Responses deviating more than two standard deviations from mean RT per participant and condition were treated as outliers, and responses to items eliciting more than 35% errors were excluded from analyses. We submitted mean RTs and accuracy data to repeated measures analyses of variance (rmANOVA) with factors SOA (100 vs. 700), Lexicality (words vs. pseudowords), and ON (many vs. few). RT results show a significant three-way interaction between SOA, lexicality, and ON, *F1* (1, 23) = 6.14, *p* < .05, *F2* (1, 844) = 8.30, *p* < .01. This three-way interaction reflects a decrease of the influence of ON with high task overlap (SOA 100); this influence, most pronounced at long SOAs, was of opposite direction for words and pseudowords: For word stimuli there was facilitation for words with many ON at the long SOA, *F1* (1, 23) = 4.24, *p* < .05 (one-tailed), *F2* (1, 418) = 7.64, *p* < .05, but not at the short SOA, F1 < 1, F2 < 1. Conversely, the impairment for pseudoword stimuli with many ON was less pronounced at short, *F1* (1, 23) = 19.71, *p* < .001, *F2* (1, 426) = 12.87, *p* < .001, than at long SOA, *F1* (1, 23) = 53.18, *p* < .0001, *F2* (1, 426) = 116.39, *p* < .0001. In other words, the typical opposite ON effects for words and pseudowords were present at long SOA but diminished or eliminated at short SOA. However, please note that the interaction between SOA and ON was significant only for pseudowords, *F1*(1, 23) = 7.40, *p* = .012, *F2*(1, 426) = 7.88, *p* = .0052, not for words, *F1*(1, 23) = .94, *p* = .34, *F2*(1, 426) = 1.56, *p* = .21. Furthermore, as expected, participants responded generally faster in conditions of low task overlap (SOA 700) than when task overlap was high (SOA 100), *F1* (1, 23) = 739.50, *p* < .0001, *F2* (1, 844) = 8611.87, *p* < .0001. There was a main effect of lexicality, with faster decisions for words than pseudowords, *F1* (1, 23) = 12.86, *p* < .01, *F2* (1, 844) = 75.54, *p* < .0001, and this effect did not interact with SOA, *F1* < 1, *F2* < 1, replicating previous findings of Rabovsky et al. (2008). We also obtained a main effect of ON, *F1* (1, 23) = 41.56, *p* < .0001, *F2* (1, 844) = 29.10, *p* < .0001, and a typical interaction between lexicality and ON, *F1* (1, 23) = 27.90, *p* < .0001, *F2* (1, 844) = 47.46, *p* < .0001.

Accuracy data for lexical decisions are presented in Table 1. Participants responded more accurately in conditions of low task overlap (SOA 700) than when task overlap was high (SOA 100), *F1* (1, 23) = 18.10, *p* < .001, *F2* (1, 844) = 26.98, *p* < .0001. Furthermore, there was a main effect of ON, *F1* (1, 23) = 17.55, *p* < .001, *F2* (1, 844) = 13.35, *p* < .001, qualified by an interaction between ON and lexicality, *F1* (1, 23) = 9.96, *p* < .01, *F2* (1, 844) = 6.06, *p* < .05. Post hoc tests showed that pseudowords with many ONs elicited more errors, *F1* (1, 23) = 18.47, *p* < .001, *F2* (1, 426) = 18.90, *p* < .0001, whereas there was no effect of ON on accuracy for words, *F1* (1, 23) = 2.54, *p* = .12, *F2* < 1. Furthermore, the three-way interaction between SOA, lexicality and ON was a trend for subjects, *F1* (1, 23) = 3.33, *p* = .081, and significant for items, *F2* (1, 844) = 4.09, *p* < .05. Post hoc tests indicated that the interaction between lexicality and ON was significant in the long SOA condition, *F1* (1, 23) = 10.20, *p* < .01, *F2* (1, 844) = 10.40, *p* < .01, but not in the short SOA condition, *F1* < 1, *F2* < 1.

**Table 1.** Performance in the tone discrimination task (RT1 and %E1), and error rates in the lexical decision (%E2)

|  | SOA short (100 ms) |  |  | SOA long (700 ms) |  |  |
| --- | --- | --- | --- | --- | --- | --- |
|  | RT1 (SEM) | %E1 (SEM) | %E2 (SEM) | RT (SEM) | %E (SEM) | %E2 (SEM) |
| Words, many ON | 777 (27) | .040 (.004) | .080 (.008) | 769 (33) | .020 (.003) | .075 (.006) |
| Words, few ON | 791 (27) | .041 (.005) | .071 (.029) | 791 (35) | .022 (.004) | .080 (.007) |
| PW, many ON | 787 (28) | .046 (.007) | .090 (.010) | 792 (35) | .025 (.005) | .081 (.009) |
| PW, few ON | 783 (28) | .051 (.006) | .064 (.006) | 775 (29) | .025 (.003) | .041 (.004) |

Note. SOA, stimulus onset asynchrony; ON, orthographic neighbours; RT, response time; E, error; SEM, standard error; PW, pseudowords.

#### Task 1

Mean correct RTs and accuracy for the tone discrimination task are presented in Table 1. We performed analogous analyses on Task 1 data as described above for the Task 2 data. Results did not show any influence of task overlap on RT1: Neither the main effect of SOA, *F* < 1, nor any interaction involving SOA reached significance, all *F*s < 1.27, *p*s > .27. The only significant effect in the RT1 analysis was an interaction between lexicality and ON, *F*(1, 23) = 6.88, *p* < .05. Follow-up tests showed a significant ON effect for words, with faster RT1 for T2 word stimuli with many as compared to few ON, *F*(1, 23) = 10.83, *p* < .01, while there was no ON effect on RT1 for T2 nonword stimuli, *F*(1, 23) = 1.79, *p* = .19.

Analyses of T1 accuracy showed a significant influence of SOA, *F*(1, 23) = 46.00, p < .0001, indicating lower accuracy in the short SOA condition, and lexicality, *F*(1, 23) = 6.11, *p* < .05, reflecting higher T1 accuracy when T2 stimuli were words as compared to pseudowords. No interaction involving SOA was significant, all *F*s < 1. As the influences of T2 variables on RT1 were not modulated by SOA (i.e., were the same independent of temporal overlap between T1 and T2), we suggest that those influences are unlikely to be meaningful in the present context. Crucially, there was no influence of SOA on RT1, suggesting that in terms of timing of responses, participants gave priority to T1, as instructed. However, the influence of SOA on T1 accuracy suggests that there was some interference of T2-related processes with T1 processing.

### 2.2 Electrophysiology

ERP effects of ON as a function of task overlap are shown in Figures 3 (for words) and 4 (for pseudowords). *F*-values and significance levels for four subsequent 100 ms time windows from 300 to 700 ms are given in Table 2. In considering the rather complex pattern of results, it seems useful to first consider the results obtained in the “SOA 700” (quotation marks indicate bold header in the table) condition, which constitutes the baseline condition. There, we clearly replicated the previously reported effects of lexicality (*F*s > 4.26 in all time segments) and orthographic neighbourhood size (*F*s > 2.58 in all time segments). How was the ON effect influenced by task overlap, that is, SOA? The results from the “Overall” analysis for the interaction between SOA and ON (E × ON × SOA) show that SOA significantly influenced ON effects in the time segment from 400 to 500 ms, and there was a trend for an influence between 300 and 400 ms. Indeed, in the “SOA 100” condition, the ON effect was not significant anymore in the segment between 400 and 500 ms, and was just a trend between 300 and 400 ms (with the corresponding effect not resembling the ON effect from the long SOA condition or previously observed ON effects; see Fig. 3 & 4). Furthermore, ON effects were also not significant in the time segments from 500 to 700 ms in the “SOA 100” condition, which contrasts with the “SOA 700” condition (even though this difference between SOAs was not reliable in the interaction analysis). Thus, ON effects on ERPs were strongly influenced by SOA, with a replication of the well-established increased N400 for words with many ON in the long SOA condition, and a lack of this effect in the short SOA condition.

**Table 2.** ANOVA results of ERP amplitudes.

| Source | df | Time segments (in ms) |  |  |  |
| --- | --- | --- | --- | --- | --- |
|  |  | 300-400 | 400-500 | 500-600 | 600-700 |
| Overall |  |  |  |  |  |
| E x SOA | 61, 1403 | 42.52*** | 63.92*** | 68.82*** | 61.23*** |
| E x LEX | 61, 1403 | 7.79*** | 16.38*** | 10.90*** | 6.23*** |
| E x ON | 61, 1403 | 2.44* | 2.25(*) | 2.57* |  |
| E x ON x SOA | 61, 1403 | 1.90(*) | 3.28** |  |  |
| E x ON x LEX | 61, 1403 | 2.44* |  |  |  |
| E x SOA x LEX | 61, 1403 |  | 4.92* | 2.08(*) |  |
| E x ON x SOA x LX | 61, 1403 |  |  |  |  |
| SOA 700 |  |  |  |  |  |
| E x LEX | 61, 1403 | 5.68*** | 10.84*** | 8.16*** | 4.27** |
| E x ON | 61, 1403 | 2.59* | 3.71** | 3.23** | 2.61* |
| E x LEX x ON | 61, 1403 | 2.02(*) |  |  |  |
| SOA 100 |  |  |  |  |  |
| E x LEX | 61, 1403 | 4.93* | 11.99*** | 6.92*** | 4.85*** |
| E x ON | 61, 1403 | 1.87(*) |  |  |  |
| Words |  |  |  |  |  |
| E x ON | 61, 1403 | 2.87* | 3.50** | 2.78* |  |
| E x ON x SOA | 61, 1403 | 1.85(*) | 1.96(*) |  |  |
| Pseudos |  |  |  |  |  |
| E x ON | 61, 1403 | 1.92(*) |  |  |  |
| E x ON x SOA | 61, 1403 |  | 2.03(*) |  |  |
| Words SOA 700 E x ON | 61, 1403 | 3.13* | 3.63** | 2.22* |  |
| Pseudos SOA 700 E x ON | 61, 1403 |  |  | 1.99(*) | 2.27* |
| Words SOA 100 E x ON | 61, 1403 |  | 1.99(*) |  |  |
| Pseudos SOA 100 E x ON | 61, 1403 |  |  |  |  |

\*\*\* $p < .001$ ; \*\* $p < .01$ ; \* $p < .05$ ; (\*) $p < .10$ . Most sources that never reached significance are omitted for the sake of clarity.

Did this pattern differ between words and pseudowords? The interaction between SOA, ON, and lexicality (E × ON × SOA × LEX) was not significant in any time segment, suggesting that the interaction between SOA and ON was not distinguishable between words and pseudowords. However, there was a significant interaction between ON and lexicality in the “Overall” analysis between 300 and 400 ms (E × ON × LEX). Following up on this interaction by considering words and pseudowords separately, revealed a significant ON effect for words and a marginally significant ON effect for pseudowords. A look at the figures reveals that the marginally significant ON effect for pseudowords between 300 and 400 ms is not just weaker than the effect for words, but different. This is unexpected as most previous studies found similar ON effects in words and pseudowords (e.g., Laszlo & Federmeier, 2011). Going beyond the significant interaction between ON and LEX in the segment between 300 and 400 ms, it seems that in general, the ON effects and their interactions with SOA were more reliable for words (compare “Words” and “Pseudos” in Table 2). However, we note that the most reliable effects were obtained when considering words and pseudowords together; therefore, we do not take this pattern to suggest fundamental differences between words and pseudowords in terms of ON-related processing, which would be at odds with previous literature, and would argue the null hypothesis. Instead, we suggest that this pattern is most plausibly explained by the overall challenge to record clean ERP data in the noisy dual task environment, requiring more trials (as provided by the joint consideration of word and pseudowords trials) for clean signals and reliable effects.

**Figure 2.**
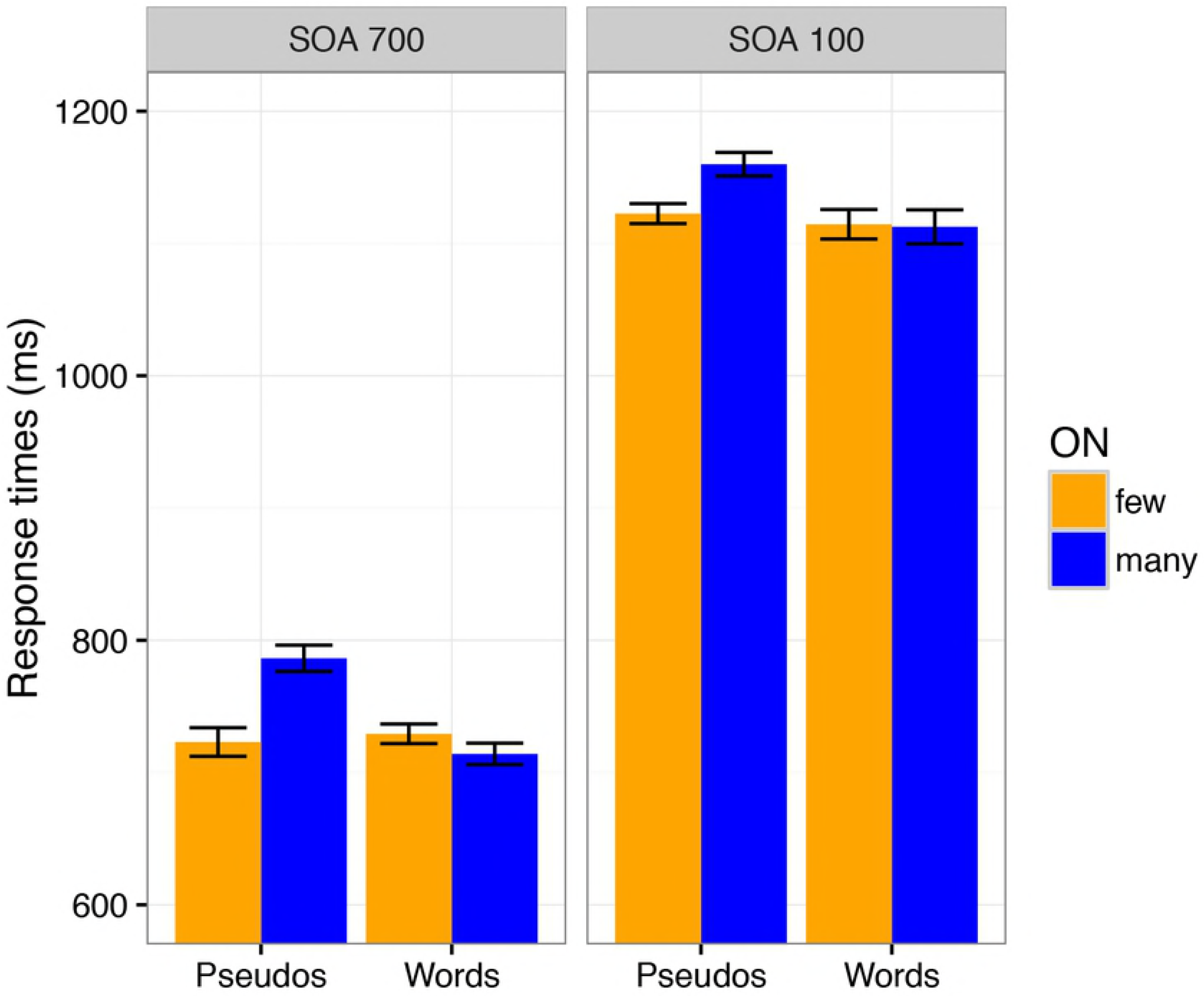
Response times as a function of stimulus onset asynchrony (SOA), lexicality (words versus pseudowords) and orthographic neighborhood size (ON). Error bars show SEMs after removal of the irrelevant overall between-subject differences (based on Cousineau, 2005).

**Figure 3.**
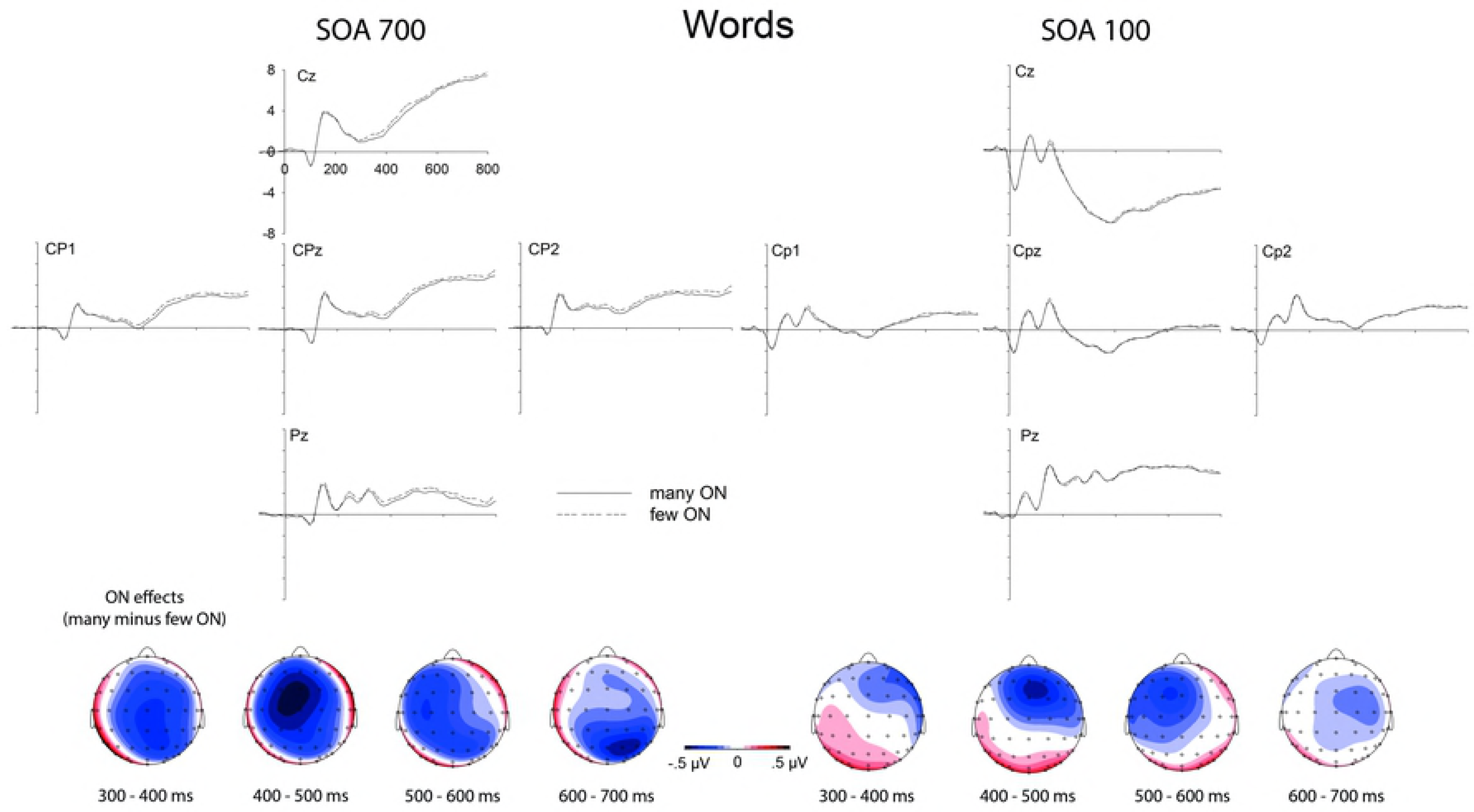
Orthographic neighborhood effects on event-related brain potentials to words. Top. Influences of orthographic neighborhood (ON) size on event-related brain potentials at centro-parietal electrode sites, to the left in conditions of low task overlap (SOA 700) and to the right in conditions of high task overlap (SOA 100). Bottom. Topographical distributions of the above depicted ON effects (many minus few ON.

**Figure 4.**
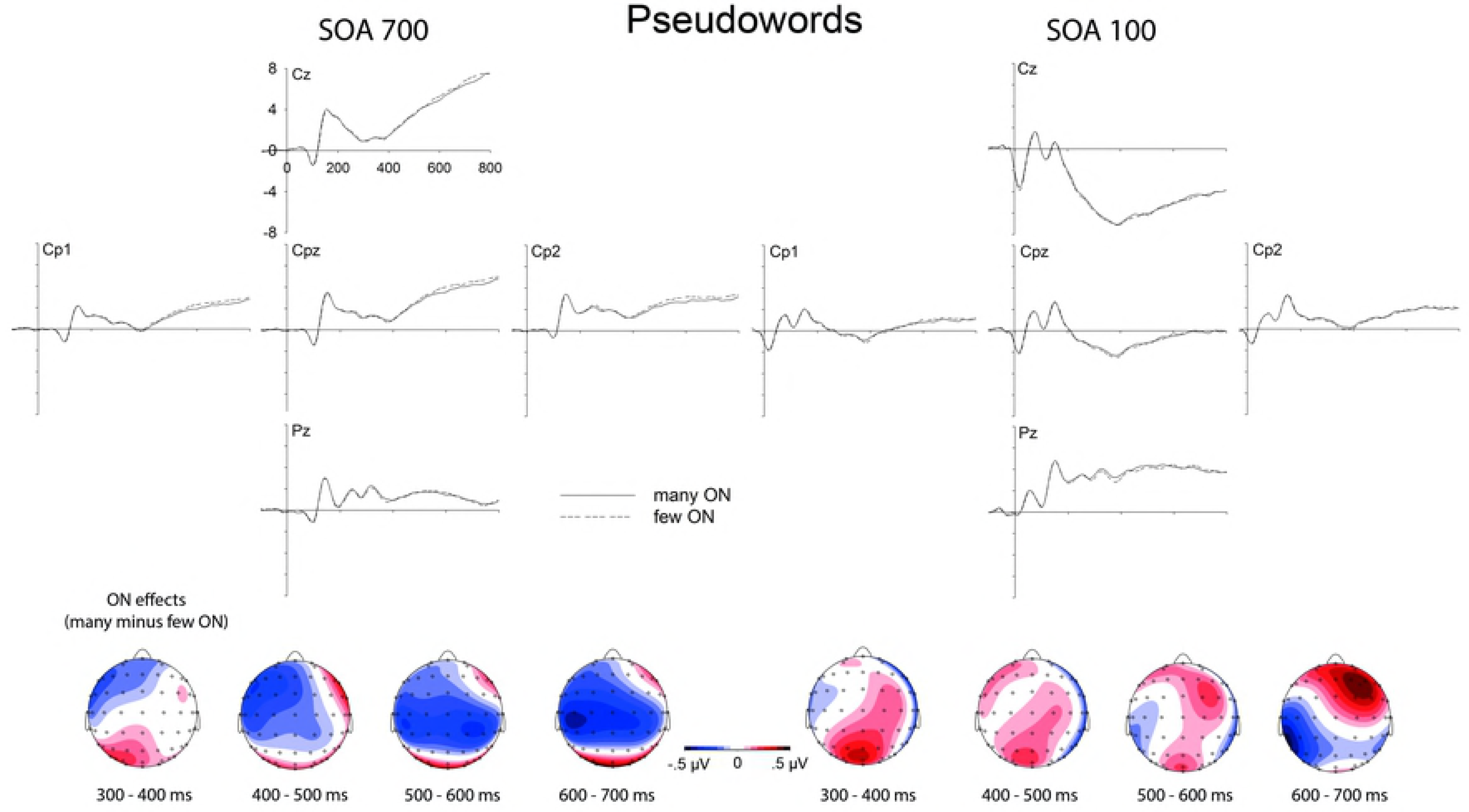
Orthographic neighborhood effects on event-related brain potentials to pseudowords. Top. Influences of orthographic neighborhood (ON) size on event-related brain potentials at centro-parietal electrode sites, to the left in conditions of low task overlap (SOA 700) and to the right in conditions of high task overlap (SOA 100). Bottom. Topographical distributions of the above depicted ON effects (many minus few ON

Significant effects of lexicality generally indicate more negativity for pseudowords than for words. Mean peak latency of the lexicality effect was earlier in the long SOA condition (492 ms) than in the short SOA condition (630 ms).

### 2.3 Comparing influences of SOA on ON effects between performance and ERPs

Here we aimed to compare the influences of SOA on ON effects in RTs relative to ERPs (see Fig. 5). This comparison is not trivial and required a number of assumptions; it intends to offer a simple perspective on the rather complex pattern of results for the interested reader. Please note that the following analyses do not aim at any scientific proof or refutation but serve to provide a certain perspective on the results. First, we re-coded the ON effects for words into the same direction as ON effects for pseudowords, under the assumption that the inverse effects for both stimulus types reflect the same underlying process, namely enhanced activation in the lexical-semantic system, facilitating the acceptance of words and impairing the rejection of pseudowords. We then compared the influence of SOA on the ON effects in words and in pseudowords (“ON effect at SOA 700” minus “ON effect at SOA 100”). In this comparison the influence of SOA on ON effects in words was not significantly different from that on pseudowords, *t*(23) = .72, *p* = .48. In order to obtain a single measure of ON effects in RTs, we averaged the ON effects on RT for words and pseudowords, separately for each SOA.

Similarly, we derived a single measure for influences of SOA on ON effects in ERPs. The global field power (GFP; Lehmann & Skrandies, 1980), reflecting the overall activity across all electrodes was averaged across the entire time segment from 300 to 700 ms. When we compared the influences of SOA on the ON effects for words and pseudowords on the GFP in the ERP, they did not differ either, *t*(23) = 1.58, *p* = .13. Hence, we also derived a single measure of the influence of SOA on ERP amplitude at each SOA.

A direct comparison of ON effects for long versus short SOA was significant, both for RTs, *t*(23) = 2.48, *p* = .021, Cohen’s *d* = .51, *M* of numerical ON effect = 20 ms, the 95% confidence interval (CI) of the numerical effect was 3 to 36 ms, i.e., 95% CI [3 ms, 36 ms], as well as ERPs, *t*(23) = 2.09, *p* = .048, Cohen’s *d* = .43, *M* = .14 μV, 95% CI [.001 μV, .287μV]. By computing the % reduction in ON effects from the long to the short SOA condition we derived a common metric of SOA-based modulations on the ON effects in behaviour and ERPs. Thus, for RTs the mean reduction was 50% with 95% CI [8%, 92%], for ERP amplitudes the mean reduction was 103% with 95% CI [1%, 206%]. Figure 5 displays effects of SOA on ON effects in RTs and ERPs side-by-side. This figure visualizes a pattern, which seems also suggested by the analyses: The reduction of the effect in ERPs seems numerically stronger (to the point that the ON effect is not significant anymore at short SOA), but the ERP data are also noisier, reducing the standardized effect size of the SOA effect.

**Figure 5.**
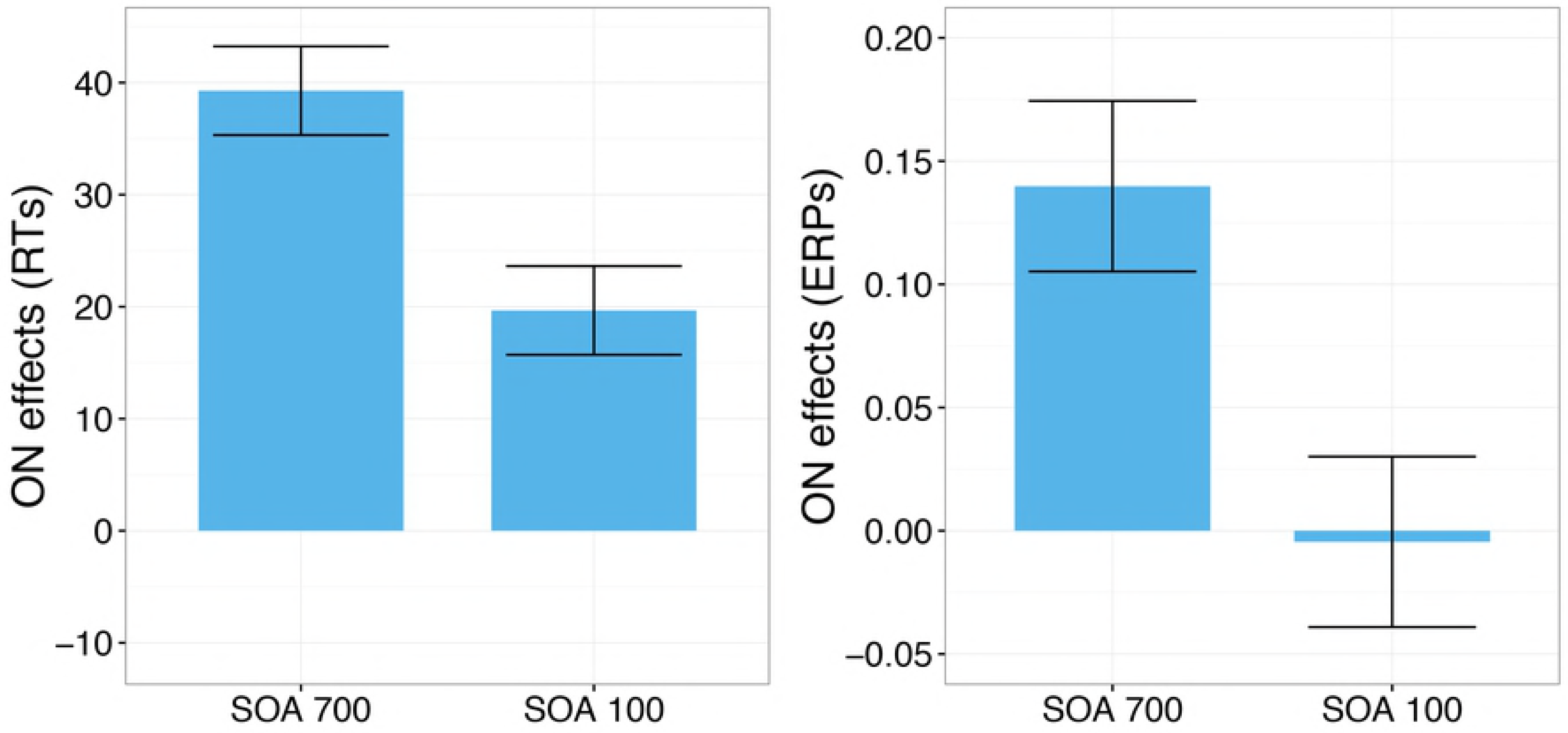
Orthographic neighbourhood effects on RTs (left) and ERPs (right) as a function of SOA (see text for details).

## 3 Discussion

The present study investigated attentional modulations of lexical processing, focusing on influences of orthographic neighbourhood size (ON) by combining a PRP paradigm with ERP recordings during lexical decisions on written words. Behavioral results show an underadditive interaction between ON and SOA (significant for pseudowords but not for words alone; see Fig. 2 and Results section). Following standard cognitive slack logic (see Fig. 1 and section 1.1 and 1.2; McCann & Johnston, 1992; Schweickert, 1978) the results would be taken to indicate that ON-related lexical processing is partly automatic and does not completely depend on central attention. At the same time, the typical N400 enhancement for stimuli with many ON which has been obtained in single task contexts (Holcomb et al., 2002; Laszlo & Federmeier, 2011), unambiguously replicated in conditions of low task overlap (SOA 700), was absent when task overlap was high (SOA 100; please see Table 2 and Fig. 3 & 4). These ERP results show that the incidental processing of orthographic neighbours during word reading is not automatic. Thus, at first sight there seems to be an incompatibility between RT and ERP data, suggesting that the application of the standard cognitive slack logic may not be appropriate for our data (see also Besner, Reynolds, & O’Malley, 2009, and below). On the other hand, the reduction in the ON effect and thus ON-related lexical processing might be seen as similar for both behaviour and ERPs (see Results section “Comparing influences of SOA on ON effects between performance and ERPs”, for details). How can we understand the observed pattern of results?

The absence of the typical ON effect on ERPs in conditions of high task overlap seems in line with a behavioural PRP study by Reynolds and Besner (2006) who investigated ON effects during reading aloud (which are facilitating in single task contexts as well; Carreiras et al., 1997; Sears et al., 1995), and reported additive effects of SOA and ON, suggesting that ON processing cannot take place when attention is focused on another task. Both this study and our ERP results suggest that the usual processing of ONs cannot take place when attention is not available. Why, then, did we observe a reduction of ON effects on lexical decision performance in the short SOA condition, that is an underadditive interaction with SOA in behavioural results, which is usually taken to indicate automaticity of the underlying processes (see section 1.1 in the introduction)? We suggest that this pattern of results was obtained because ON-related processes are not necessary for lexical decisions and can be omitted or reduced. This suggestion highlights an implicit assumption necessary for the application of cognitive slack logic, namely that the process of interest is necessary for task performance (see further discussion in section ‘Logic of cognitive slack, partial underadditivity, and limitations’, below). We detail this perspective in the following and discuss two specific possible interpretations differing in their assumptions concerning which aspect of ON-related processing is omitted or reduced when central attention is unavailable.

### 3.1 The activation of orthographic neighbours may be omitted or reduced

First, ON activation may not be necessary for lexical decisions. Reynolds and Besner (2006) who observed an additive interaction between SOA and ON in reading aloud (see above) took ON effects in their study to reflect lexical contributions to phonological recoding. These contributions were apparently necessary for the task of reading aloud and had to be made up for when attention was available again, resulting in additive effects of SOA and ON. Even though phonological processing has been shown to be involved in word processing under normal reading conditions (see, e.g., Conrad et al., 2007; Ziegler & Jacobs, 1995; Ziegler et al., 2000), it is not clear whether it is strictly necessary for deciding whether a stimulus is a word or not, and ON activation may not be as essential for lexical decisions as they are for reading aloud. Thus, in the present LDT experiment, lexical processing supporting lexical decisions may have taken place during the slack (i.e. while central attention was unavailable) despite impeded ON activation. Therefore, there might have been no need to make up for the initial lack of ON activation when attention was eventually focused on the lexical decision.

From this perspective, the underadditive interaction observed in the present study would not indicate automaticity of ON activation (i.e. taking place during slack) but would rather suggest its absence, that is, lexical processing taking place without or with reduced ON activation during the slack. Lexical processing during the slack period is also suggested by lexicality effects on the ERP – often seen as electrophysiological indicators of lexical processing (Braun et al., 2006; Hutzler, Bergmann et al., 2004) – being present in the short SOA condition and only slightly delayed as compared to the long SOA condition. Specifically, the mean peak latency of the lexicality effect was 492 ms in the long SOA condition and 630 ms in the short SOA condition. These findings replicate the slight delay of lexical processing under conditions of high task overlap observed in our previous study (Rabovsky et al., 2008); that is, in both cases the delay of lexicality effects in the ERPs (peak latency difference of 138 ms in the current study and 104 ms in the study by Rabovsky et al., 2008) was smaller than the delay in behavioural responses (384 ms in the current study and 385 ms in Rabovsky et al., 2008), and hence smaller than what would be expected when the underlying processes had been postponed until attention was available again because in this case the delay in ERP effects would be expected to be equally large as the delay in behavioural responses (Arnell et al., 2004; Luck, 1998). Lexicality effects presumably reflect the difference in brain activity between a situation in which the system settles into an existing lexical semantic representation (for words) or not (for pseudowords). This settling process was apparently slowed but not precluded by task overlap.

A possible explanation for both the slight delay of lexical processing and the lack of ON activation might be that participants inhibit the lexical-semantic system under conditions of high task overlap in order to optimize T1 performance via top-down attentional control (De Jong, 1995; Rabovsky et al., 2008). Even within an inhibited system, the target words may eventually (somewhat later than usual) be recognized, or cross their activation threshold in terms of computational models of visual word recognition, because their representations receive continuous support from visual input. However, the lexical and semantic representations of the orthographic neighbours that receive less input activation from letter representations than representations of the target words (because they do not match all aspects of the orthographic input) may not become sufficiently activated when the lexical-semantic system is inhibited, resulting in the absence of the typical N400 enhancement for stimuli with a large orthographic neighbourhood which we observed in the short SOA condition.

### 3.2 The inhibition of orthographic neighbours may be omitted or reduced

A second possible interpretation of the observed pattern of results is based on the assumption that the influence of ON size on N400 amplitudes does not reflect the activation of the lexical and/ or semantic representations of the orthographic neighbours (Braun et al., 2006; Holcomb et al., 2002; Laszlo & Federmeier, 2011; Laszlo & Plaut, 2012; Müller et al., 2010) but – to the contrary – their inhibition (Debruille, 1998, 2007; Molinaro et al., 2010). In this case, though activation of ONs may proceed while attention is unavailable, their inhibition may depend on central attention and may therefore be reduced when attention is focused on another task - resulting in the absence of the typical ON effect on N400 amplitudes in the short SOA condition. It is important to note that, different from the top-down inhibition discussed in the previous section, the inhibition process assumed here is based on lateral inhibition between neighbouring representations. Even though pseudowords do not have lexical representations, the lexical representations of their orthographic neighbours may be activated incidentally yielding inhibition between these co-activated representations.

The need to inhibit ONs for target identification is apparent in results showing that large ONs may also slow down visual word recognition, especially in tasks requiring unambiguous identification and complete lexical access rather than a mere “word” response or a pronunciation, for example, in perceptual identification tasks (e.g. Carreiras et al., 1997; for a review see Andrews, 1997). Opposing influences of ON size have been simulated in a model of visual word recognition (Grainger & Jacobs, 1996), assuming that “word” responses in lexical decision tasks can be given either via a fast-guess criterion based on global lexical activation, yielding facilitation by ON size, or via an identification criterion based on complete lexical access requiring more inhibition as ON size increases. In our PRP paradigm, fast-guess responses and thus facilitation by larger ON size are only possible in the long SOA condition, where we indeed obtained a facilitating influence of increasing ON size. In contrast, in the short SOA condition, central attention necessary for decision making is not available early on so that fast-guess responses are precluded by the task context. Later on, when central attention is available for the lexical decision, lexical access to the target will already have taken place, so that ON size should have less of an influence, as indeed observed in the short SOA condition. While the inhibition of ONs is involved in lexical processing under normal conditions where it allows for unambiguous identification of the target, it may not always be strictly necessary: Even when neighbour representations are not (or less) actively inhibited, the longer the processing of the target proceeds, the more will target representations dominate all other word representations because they are the only ones that completely match the visual input and therefore receive the highest amount of activation from letter representations. In this view, even with high task overlap, lexical access to the target word could apparently take place, albeit with a slight delay (as indicated by the ERP lexicality effect) and in the absence of neighbour inhibition (as indicated by the absence of the typical ON effect on N400 amplitudes).

### 3.3 Logic of cognitive slack, partial underadditivity, and limitations

Finally, as noted above our results may have implications for the interpretation of underadditivity in the PRP paradigm more generally. Following standard cognitive slack logic (see introduction and Fig. 1), an underadditive interaction of the influences of a difficulty manipulation of a given T2 related process and SOA is taken to suggest automaticity of this T2 related process. Our results can be seen as suggesting that this is not always the case and that such an interpretation depends on further assumptions such as the assumption that the respective process is necessary for task performance. The general conclusion that there are exceptions to cognitive slack logic is in line with a study by Besner, Reynolds, and O’Malley (2009) demonstrating that cognitive slack logic does not apply when Task 2 involves two competing processes that operate in parallel, and Task 1 interferes with only one of them. Together, both studies show that cognitive slack logic depends on additional assumptions and careful consideration of the functional architecture of the processes under investigation is necessary to draw conclusions concerning their automaticity.

It is important to note that what we observed was a pattern of partial underadditivity, because the ON effect was reduced, but not eliminated, in the short SOA condition (with an estimated 50% reduction; see Fig. 5). Therefore, ON-related processes did occur after the slack but to a smaller degree than when central attention was immediately available, as in the long SOA condition, and also less than to be expected when the relevant process had been merely postponed until after the slack. Therefore, the discussion above needs to be qualified to reflect the observation that ON-related processing (co-activation or inhibition) was not completely omitted when attention was initially unavailable but merely reduced. Together with the absence of an ERP effect in the short SOA condition (with an estimated 100% reduction), this may suggest that while attention is unavailable, ON-related processing does not take place but that only part of it (estimated 50%) needs to be made up for, or in any case, according to our data, *is* made up for when attention is available again.

An interesting question in this context is why the reduction of the ON effect by SOA appeared to differ between RTs and ERPs, and why the apparent influence that ON had on processing after the slack was not visible in ERPs. One possible answer lies in the need for time-locking to relevant events to observe reliable ERP effects. We time-locked the ERPs with respect to the presentation of the word/pseudoword stimuli, but the ON related processes after the slack may not have been tightly time-locked with respect to that event, and may instead have been synchronized to the clearance of the bottleneck, which may vary to some extent across trials, leading to smearing of the ERPs during averaging, which diminishes the size of conditions differences (Ouyang, Sommer & Zhou, 2016). Another possibility might be that ON influences not only lexical processing, reflected in N400 amplitudes, but also decision processes, which may not be reflected in N400 amplitudes, and that the former but not the latter aspects are omitted under conditions of high task overlap. Further research is required to investigate these issues.

A potential limitation of our study is that the tone discrimination task was relatively difficult (800 Hz versus 1000 Hz) and it is not entirely clear whether results would be the same with an easier discrimination (e.g., 200 Hz versus 1000 Hz). An important issue in PRP studies is whether the interference is functionally localized at the level of central attention (e.g., decision making) or input attention (i.e., perceptual processes). In designing the study, we reasoned that an auditory tone task would not interfere with a visual word recognition task at the level of input attention, with interference instead being limited to the central stage of decision and response selection. Nonetheless, phonological processes may be incidentally involved in visual word recognition (see, e.g., Conrad et al., 2007; Ziegler & Jacobs, 1995; Ziegler et al., 2000) and thus it seems in principle possible that the perceptual processes involved in task 1 interfered with phonological processes occurring during the visual lexical decision task 2. Future research using an easier auditory discrimination task should investigate this possibility.

## 4 Conclusions

In sum, combining the PRP paradigm with ERPs we found that the typical N400 enhancement for stimuli with many ONs during lexical decisions on written words was absent under conditions of high task overlap. Together with the observation that ERP lexicality effects, assumed to reflect lexical processing supporting lexical decisions, were somewhat delayed but present even in conditions of high task overlap, the absence of the typical ON effect on N400 amplitudes indicates that at least some of the processes induced by ON may not be essential for lexical processing. These findings show that the incidental processing of orthographic neighbors during word reading is not automatic.

## 5 Experimental Procedure

### 5.1 Ethics statement

This research has been conducted according to the principles expressed in the Declaration of Helsinki and was approved by the Ethics Committee at the Department of Psychology, University of La Laguna, Tenerife. Participants provided written informed consent prior to participation.

### 5.2 Participants

Twenty-four students (14 women and 10 men) of the University of La Laguna, Tenerife, with a mean age of 23 years (ranging between 18 and 31) participated in the experiment. All of them were native Spanish speakers. According to the Edinburgh Handedness Inventory (Oldfield, 1971), 22 of the participants were right-handed and 2 were ambidextrous. They either received course credits or monetary compensation for participation.

### 5.3 Stimuli and apparatus

Stimuli used for T2 were 480 bisyllabic Spanish words, selected from the LEXESP Spanish database (Sebastián-Galles et al., 2000), and 480 pseudowords. Pseudowords were constructed by exchanging first and second syllables of word stimuli pertaining to the same experimental condition. All pseudowords were pronounceable, meeting overall characteristics and syllabic structure of word stimuli, and all words were content words. We selected stimuli such that the number of orthographic neighbours differed for both words (*M*s = 13.0 vs. 2.9) and pseudowords (*M*s = 11.5 vs. 1.8). Stimuli (words and nonwords) were entered in the high ON condition when they had more than 5 orthographic neighbours and stimuli with no more than 5 orthographic neighbours were entered in the low ON condition. All stimuli had between 4-6 letters assuring for foveal processing. Exact stimulus length was controlled for using almost identical numbers of stimuli of specific length across conditions (51-55 four letter words, 161-169 five letter words, and 20-25 six letter words across the four stimulus conditions; *M*s = 4.9 in all conditions). Frequency of stimulus words was spreading naturally across the range of word frequency from 1-300 per 1 Million of Occurrences in the CELEX database. The logarithm (base 10) of this Word Frequency was held constant across conditions of word stimuli (*M*s = 1.0 for the LOG(10) of word frequency in both conditions) with about half of the stimuli in each condition being of low word frequency (<10 per 1 Million of occurrences). Phonological neighbourhood size (PN) was not controlled for; the correlation between ON and PN within the word stimuli was *r* = .79. A high correlation between ON and PN is to be expected for Spanish stimuli, as Spanish is a language with a very high consistency between orthography and phonology. For reading aloud, it has been shown that when ON is confounded with PN, the effect occurs at phonology (e.g., Mulatti, Reynolds, & Besner, 2006), but for lexical decisions the pattern is more complex, with inverse effects of PN for high versus low ON words in both behavioral performance and N400 amplitudes (Carrasco-Ortiz, Midgley, Grainger, & Holcomb, 2017; Grainger, Muneaux, Farioli, & Ziegler, 2005). These effects have been obtained with English stimuli and have been interpreted in terms of cross-code consistency, that is, due to the compatibility of orthographic and phonological representations. Because this compatibility is generally very high for Spanish words, further research is required to investigate in how far these results generalize to Spanish. For current purposes, we note that we cannot differentially attribute our effects to either ON or PN. The stimuli were presented in light grey lower case letters in the middle of a dark grey screen, positioned at eye level 1 m in front of the participants.

Stimuli for T1 were two sinusoidal tones of 800 and 1000 Hz and 60 ms duration presented via two loudspeakers located to the left and right of the screen. For the manual responses two marked keys of a standard keyboard were operated with the index fingers. Foot responses were recorded with two USB mice fixated on a wooden footrest to be easily reached and pressed with the big toes, shoes being taken off. Stimulus presentation and recording of responses was controlled by Presentation software (Neurobehavioral Systems, Albany, CA, USA).

### 5.4 Procedure

The trial scheme is depicted in Figure 6. A trial started with a fixation point at the middle of the screen. After an interval of 2 s one of the tones was presented. Participants were to indicate as fast and accurately as possible with their left or right foot whether the tone was of high or low pitch. The LDT stimuli were presented 100 or 700 ms after tone onset. Participants were instructed to press the key labeled “Sí” (yes) if the letter string was a real Spanish word or to press the key labeled “No” (no) if it was not. Keys were to be pressed with the left or right index finger, with assignment of response hand to response type counterbalanced across participants. Participants were instructed to perform the LDT as fast and accurately as possible, but to give priority to the pitch discrimination task. The letter string remained on the screen until both responses had been given or until 2.5 s had elapsed.

**Figure 6.**
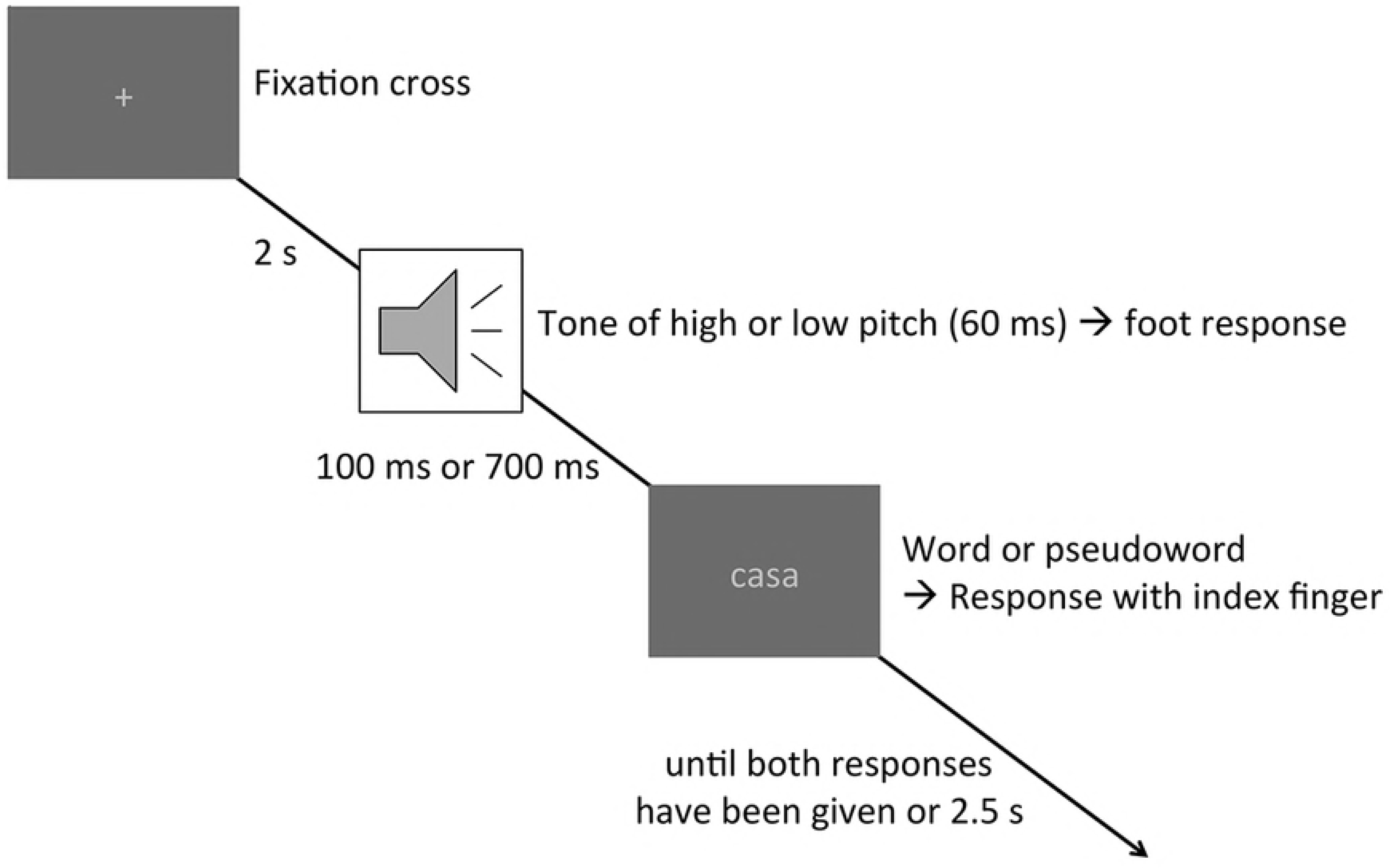
Trial scheme. A trial started with a fixation cross. After 2 s, a tone of 60 ms duration and either high or low pitch was presented, requiring a pitch discrimination via a foot response (task 1). After a stimulus onset asynchrony (SOA) of either 100 or 700 ms, a letter string (either a word or a pseudoword) replaced the fixation cross on the screen, requiring a lexical decision via button press with the left or right index finger (Task 2). The letter string remained on the screen until both responses had been given for a maximum of 2.5 ms.

The next experimental trial started immediately thereafter or following a written feedback, presented for 500 ms on the screen. Error feedback was given if one of the key presses was incorrect or missing or if the LDT response (R2) had been given before R1.

There were three practice blocks of 48 trials each to familiarize participants with the task requirements. The pitch discrimination task and LDT were first practiced in single-task blocks, and then in combination in one dual-task block. For practice trials, different letter strings were used than in the experiment proper. The experiment consisted of 960 trials, subdivided into 16 blocks of sixty trials each with a pause after each block. All possible condition combinations of tone pitch, SOA, lexicality, and ON were of equal probability, and each letter string appeared only once. The assignment of the words to the long and short SOA conditions was counterbalanced such that overall, each word appeared equally often in both conditions. Trials appeared in a different random order for each participant so that the different SOAs were presented intermixed. The assignments of tone pitch to response foot and of lexicality to response hand were counterbalanced across participants.

### 5.5 EEG recording

The EEG was recorded from 64 Ag/AgCl-electrodes mounted in a Quik-Cap electrode cap (Compumedics Neuroscan, Charlotte, NC, USA) with a left mastoid reference. External electrodes were used for the reference electrode, the right mastoid, and the horizontal and vertical electrooculograms, which were recorded bipolarly from the external canthi and from above and below the left eye. Conductive Quik-Gel (Compumedics Neuromedical Supplies, Charlotte, NC, USA) was used as electrolyte. Signals were recorded with a SynAmps2 amplifier (Compumedics Neuroscan, Charlotte, NC, USA), set to a bandpass of 0.032–70 Hz and a 50 Hz notch filter; sampling rate was 500 Hz.

Offline, the continuous EEG was segmented into epochs of 1800 ms, starting 800 ms before letter string onset. After applying an average reference (proposed as an estimate for an inactive reference and considered to be less biased than other common references; Picton et al., 2002) and a 30 Hz low pass filter, eye blink artifacts were removed with a spatio-temporal dipole modelling procedure using BESA software (Berg & Scherg, 1991). Trials containing other artefacts and trials with incorrect or missing responses were discarded. Before generating averages for each participant, electrode, and experimental condition, the epochs were referred to a baseline starting 100 ms prior to the LDT stimuli.

ERPs were submitted to rmANOVAs with factors Electrode Site (61 levels), SOA (100 vs. 700 ms), Lexicality (words vs. pseudowords), and ON (many vs. few). Because of the average reference, only effects in interaction with Electrode Site are meaningful in these ANOVAs and are considered as “main effects” of the respective factors. Follow-up tests consisted in subsequent rmANOVAs focusing on subsets of conditions. Analyses focused on four subsequent segments of 100 ms each, starting at 300 ms after word onset.

## Acknowledgements

This project has received funding from the European Union’s Horizon 2020 research and innovation programme under the Marie Sklodowska-Curie grant agreement No 658999 to Milena Rabovsky and by PSI2013-47959-P (Spanish Government) to the third author. The funding sources were not involved in any aspect of this research. We thank David Beltrán for excellent technical support and help during set-up and data acquisition.

## References

Allen, P. A., Lien, M.-C., Murphy, M. D., Sanders, R. E., Judge, K. S., & McCann, S. R. (2002). Age differences in overlapping-task performance: Evidence for efficient parallel processing in older adults. Psychology and Aging, 17, 505–519.

Andrews, S. (1989). Frequency and neighborhood effects on lexical access – activation or search. Journal of Experimental Psychology: Learning, Memory, and Cognition, 15, 802–814.

Andrews, S. (1997). The effect of orthographic similarity on lexical retrieval: resolving neighborhood conflicts. Psychonomic Bulletin & Review, 4, 439–461.

Arnell, K.M., Helion, A.M., Hurdelbrink, J.A., & Pasieka, B. (2004). Dissociating sources of dual-task interference using human electrophysiology. Psychonomic Bulletin & Review, 11, 77–83.

Augustinova, M & Ferrand, L. (2014). Automaticity of word reading: Evidence from the Semantic Stroop Paradigm. Current Directions in Psychological Science, 23, 343–348.

Barber, H., Vergara, M., & Carreiras, M. (2004). Syllable-frequency effects in visual word recognition: evidence from ERPs. Neuroreport, 15, 545–548.

Berg, P. & Scherg, M (1991). Dipole modeling of eye activity and its application to the removal of eye artefacts from the EEG and MEG. Clinical Physiology and Physiological Measurements, 12, 49–54.

Besner, D., Reynolds, M., & O’Malley, S. (2009). When underadditivity of factor effects in the Psychological Refractory Period paradigm implies a bottleneck: Evidence from psycholinguistics. The Quarterly Journal of Experimental Psychology, 62, 2222–2234.

Besner, D., Risko, E.F., Stolz, J.A., White, D., Reynolds, M., O’Malley, S., & Robidoux, S. (2016). Varieties of attention: Their roles in visual word identification. Current Directions in Psychological Science, 25, 162–168.

Braun, M., Jacobs, A. M., Hahne, A., Ricker, B., Hofmann, M., & Hutzler, F. (2006). Model-generated lexical activity predicts graded ERP amplitudes in lexical decision. Brain Research, 1073-1074, 431–439.

Brisson, B. & Jolicoeur, P., (2007). A psychological refractory period in access to visual short-term memory and the deployment of visual-spatial attention: multitasking processing deficits revealed by event-related potentials. Psychophysiology 44, 323–333.

Broadbent, D. E. (1967). Word-frequency effect and response bias. Psychological Review, 74, 1–15.

Carrasco-Ortiz, H., Midgley, K.J., Grainger, J., & Holcomb, P.J. (2017). Interactions in the neighborhood: Effects of orthographic and phonological neighbors on N400 amplitude. Journal of Neurolinguistics, 41, 1–10.

Carreiras, M., Perea, M., & Grainger, J. (1997). Effects of Orthographic Neighborhood in Visual Word Recognition: Cross-Task Comparisons. Journal of Experimental Psychology: Learning, Memory, and Cognition, 23(4), 857–871

Cleland, A. A., Gaskell, M. G., Quinlan, P. T., & Tamminen, J. (2006). Frequency effects in spoken and visual word recognition: Evidence from dual-task methodologies. Journal of Experimental Psychology: Human Perception and Performance, 32, 104–119.

Coltheart, M., Davelaar, E., Jonasson, J. T., & Besner, D. (1977). Access to the internal lexicon. In S. Dornic (Ed.), Attention and Performance VI (pp. 535–555). New York: Erlbaum.

Coltheart, M., Rastle, K., Perry, C., Langdon, R., & Ziegler, J. (2001). DRC: A dual route cascaded model of visual word recognition and reading aloud. Psychological Review, 108, 204–256.

Conrad, M., Grainger, J., & Jacobs, A. M. (2007). Phonology as the source of syllable frequency effects in visual word recognition: Evidence from French. Memory & Cognition, 35 (5), 974–983.

Cousineau, D. (2005). Confidence intervals in within-subject designs: A simpler solution to Loftus and Masson’s method. Tutorials in Quantitative Methods for Psychology, 1, 42–45.

Debruille, J. B. (1998). Knowledge inhibition and N400: A study with words that look like common words. Brain and Language, 62, 202–220.

Debruille, J. B. (2007). The N400 potential could index a semantic inhibition. Brain Research Reviews, 56, 472–477.

De Jong, R. (1995). The role of preparation in overlapping task performance. The Quarterly Journal of Experimental Psychology. A, Human Experimental Psychology, 48, 2–25.

Dell’Acqua, R., Jolicoeur, P., Vespignani, F., & Toffanin, P., (2005). Central processing overlap modulates P3 latency. Experimental Brain Research, 165, 54–68.

Forster, K. I. (1976). Accessing the mental lexicon. In R. J. Wales & E. W. Walker (Eds.), New approaches to language mechanisms (pp. 257–287).

Amsterdam: North-Holland. Forster, K. I., & Chambers, S. M. (1973). Lexical access and naming time. Journal of Verbal Learning and Verbal Behavior, 12, 627–635.

Forster, K.I. & Shen, D. (1996). No enemies in the neighborhood: absence of inhibitory neighborhood effects in lexical decision and semantic categorization. Journal of Experimental Psychology: Learning, Memory, & Cognition, 22, 696–713.

Frank, S.L., Otten, L.J., Galli, G., & Vigliocco, G. (2015). The ERP response to the amount of information conveyed by words in sentences. Brain and Language, 140, 1–11.

Grainger, J. & Jacobs, A.M. (1996). Orthographic processing in visual word recognition: a multiple read-out model. Psychological Review, 103, 518–565.

Grainger, J., Muneaux, M., Farioli, F., & Ziegler, J.C. (2005). Effects of phonological and orthographic neighborhood density interact in visual word recognition. The Quarterly Journal of Experimental Psychology, 58A, 981–998.

Hohlfeld, A., Mierke, K., & Sommer, W. (2004). Is word perception in a second language more vulnerable than in one’s native language? Evidence from brain potentials in a dual task setting. Brain and Language, 89, 569–579.

Hohlfeld, A., Sangals, J., & Sommer, W. (2004). Effects of additional tasks on language perception: An event-related brain potential investigation. Journal of Experimental Psychology: Learning, Memory, and Cognition, 30, 1012–1025.

Holcomb, P. J., Grainger, J. & O’Rourke, T. (2002). An electrophysiological study of the effects of orthographic neighborhood size on printed word perception. Journal of Cognitive Neuroscience, 14, 938–950.

Howes, D. H., & Solomon R. L. (1951). Visual duration threshold as a function of word-probability. Journal of Experimental Psychology, 41, 401–410.

Hutzler, F., Bergmann, J., Conrad, M., Kronbichler, M., Stenneken, P., & Jacobs, A. M. (2004). Inhibitory effects of first syllable-frequency in lexical decision: an event-related potential study. Neuroscience Letters, 372, 179–184.

Huynh, H., & Feldt, L. S. (1976). Estimation of the box correction for degrees of freedom from sample data in randomized block and split-plot designs. Journal of Educational Statistics, 1, 69–82.

Johnston, J. C., McCann, R. S., & Remington, R. W. (1995). Chronometric evidence for two types of attention. Psychological Science, 6, 365–369.

Kahnemann, D. (1973). Attention and effort. Englewood Cliffs, NJ: Prentice Hall.

Keele, S. W. (1973). Attention and human performance. Pacific Palisades, CA: Goodyear.

Kounios, J. & Holcomb, P. J. (1994). Concreteness effects in semantic processing: ERP evidence supporting dual-coding theory. Journal of Experimental Psychology: Learning, Memory, and Cognition, 20, 804–823.

Kutas, M. & Federmeier, K. D. (2011). Thirty years and counting: Finding meaning in the N400 component of the event-related brain potential (ERP). Annual Review of Psychology, 62, 14.1–14.27.

Laszlo, S. & Armstrong, B.C. (2014). PSPs and ERPs: Applying the dynamics of postsynaptic potentials to individual units in simulation of temporally extended event-related potential reading data. Brain and Language, 132, 22–27.

Lazlo, S. & Federmeier, K. D. (2011). The N400 as a snapshot of interactive processing: Evidence from regression analyses of orthographic neighbor and lexical associate effects. Psychophysiology, 48, 176–186.

Laszlo, S. & Plaut, D.C. (2012). A neurally plausible parallel distributed processing model of event-related potential word reading data. Brain and Language, 120, 271–281.

Lehmann, D., and Skrandies, W. (1980). Reference-free identification of components of checkerboard-evoked multichannel potential fields. Electroencephalography and Clinincal Neurophysioly, 48, 609–621.

Lien, M.-C., Allen, P. A., Ruthruff, E., Grabbe, J., McCann, R. S., & Remington, R. W. (2006). Visual word recognition without central attention: Evidence for greater automaticity with advancing age. Psychology and Aging, 21, 431–447.

Lien, M.-C., Ruthruff, E., Cornett, L., Goodin, Z., & Allen, P. A. (2008). On the nonautomaticity of visual word processing: electrophysiological evidence that word processing requires central attention. Journal of Experimental Psychology: Human Perception and Performance, 34, 751–773.

Luck, S. J. (1998). Sources of dual-task interference: Evidence from human electrophysiology. Psychological Science, 9, 223–227.

Martín-Loeches, M., Schacht, A., Casado, P., Hohlfeld, A., Abdel Rahman, R., & Sommer, W. (2009). Rules and heuristics during sentence comprehension: evidences from a dual-task brain potential study. Journal of Cognitive Neuroscience, 21, 1380–1395.

McCann, R. S., & Johnston, J. C. (1992). Locus of the single-channel bottleneck in dual-task interference. Journal of Experimental Psychology: Human Perception and Performance, 18, 471–484.

McCann, R. S., Remington, R. W., & Van Selst, M. (2000). A dual-task investigation of automaticity in visual word processing. Journal of Experimental Psychology: Human Perception and Performance, 26, 1352–1370.

McClelland, J. L., & Rumelhart, D. E. (1981). An interactive activation model of context effects in letter perception: Part 1. An account of basic findings. Psychological Review, 88, 375–407.

McLeod, P. (1977). Parallel processing and the psychological refractory period. Acta Psychologica, 41, 381–391.

Molinaro, N., Conrad, M., Barber, H.. & Carreiras, M. (2010). On the functional nature of the N400: Contrasting effects related to visual word recognition and contextual semantic integration. Cognitive Neuroscience, 1: 1, 1—7.

Monsell, S. (1991). The nature and locus of word frequency effects in reading. In D. Besner & G. W. Humphreys (Eds.), Basic processes in reading: Visual word recognition (pp. 148–197). Hillsdale, NJ: Erlbaum.

Monsell, S., Doyle, M. C., & Haggard, P. N. (1989). Effects of frequency on visual word recognition tasks: Where are they? Journal of Experimental Psychology: General, 118, 43–71.

Morton, J. (1969). Interaction of information in word processing. Psychological Review, 76, 165–178.

Mulatti, C., Reynolds, M., & Besner, D. (2006). Neighborhood effects in reading aloud: New findings and new challenges for computational models. Journal of Experimental Psychology: Human Perception and Performance, 32, 799–810.

Müller, O., Duñabeitia, J. A., & Carreiras, M. (2010). Orthographic and associative neighborhood density effects: What is shared, what is different? Psychophysiology, 47, 455–466.

Nuñez, P. L. (1981). Electric fields of the brain: The neurophysics of EEG. New York: Oxford University Press.

Oldfield, R. C. (1971). The assessment and analysis of handedness: The Edinburgh inventory. Neuropsychologia, 9, 97–113.

O’Malley, S., Reynolds, M.G., Stolz, J.A., & Besner, D. (2008). Reading aloud: Spelling-sound translation uses central attention. Journal of Experimental Psychology: Learning, Memory, and Cognition, 34, 422–429.

Osman, A., & Moore, C. M. (1993). The locus of dual-task interference: Psychological refractory effects on movement-related brain potentials. Journal of Experimental Psychology: Human Perception and Performance, 19, 1292–1312.

Ouyang, G., Sommer, W., & Zhou, C. (2016). Reconstructing ERP amplitude effects after compensating for trial-to-trial latency jitter: A solution based on a novel application of residue iteration decomposition. International Journal of Psychophysiology, 109, 9–20.

Pashler, H. (1984). Processing stages in overlapping tasks: Evidence for a central bottleneck. Journal of Experimental Psychology: Human perception and performance, 10, 358–377.

Pashler, H., & Johnston, J. C. (1989). Chronometric evidence for central postponement in temporally overlapping tasks. Quarterly Journal of Experimental Psychology, 41A, 19–45.

Picton, T. W., Bentin, S., Berg, P., Donchin, E., Hillyard, S. A., Johnson, R., et al. (2000). Guidelines for using human event-related potentials to study cognition: Recording standards and publication criteria. Psychophysiology, 37, 127–152.

Plaut, D. C., McClelland, J. L., Seidenberg, M. S., & Patterson, K. (1996). Understanding normal and impaired word reading: Computational principles. Psychological Review, 103, 56–115.

Rabovsky, M., Álvarez, C. J., Hohlfeld, A., & Sommer, W. (2008). Is lexical access autonomous? Evidence from combining overlapping task with recording event-related brain potentials. Brain Research, 1222, 156–165.

Rabovsky, M., Hansen, S.S., & McClelland, J.L. (2016). N400 amplitudes reflect change in a probabilistic representation of meaning: Evidence from a connectionist model. Proceedings of the 38th Annual Meeting of the Cognitive Science Society, 2045–2050. Cognitive Science Society: Austin, TX.

Rabovsky, M. & McRae, K. (2014). Simulating the N400 ERP component as semantic network error: Insights from a feature-based connectionist attractor model of word meaning. Cognition, 132, 68–89.

Rellecke, J., Bakirtas, A.M., Sommer, W., & Schacht, A. (2011). Automaticity in attractive face processing: brain potentials from a dual task. Neuroreport, 22, 706–710.

Reynolds, M & Besner, D. (2006). Reading aloud is not automatic: Processing capacity is required to generate a phonological code from print. Journal of Experimental Psychology: Learning, Memory, and Cognition, 6, 1303–1323.

Rubenstein, H., Garfield, H., & Millikan, J. A. (1970). Homographic entries in the internal lexicon. Journal of Verbal Learning and Verbal Behavior, 9, 487–494.

Ruthruff, E., Allen, P.A., Lien, M.-C., & Grabbe, J. (2008). Visual word recognition without central attention: Evidence for greater automaticity with greater reading ability. Psychonomic Bulletin & Review, 15, 337–343.

Schweickert, R. (1978). A critical path generalization of the additive factor method: Analysis of a Stroop task. Journal of Mathematical Psychology, 18, 105–139.

Sears, C.R., Hino, Y. & Lupker, S.J. (1995). Neighborhood size and neighborhood frequency effects in visual word recognition. Journal of Experimental Psychology: Human Perception and Performance, 21, 876–900.

Sebastián-Gallés, N., Martí, M. A., Carreiras, M., & Cuetos, F. (2000). LEXESP: Una base de datos informatizada del español [LEXESP: A computerized database of Spanish]. Barcelona, Spain: Universitat de Barcelona.

Seidenberg, M. S., & McClelland, J. L. (1989). A distributed developmental model of word recognition and naming. Psychological Review, 96, 523–568.

Sommer, W. & Hohlfeld, A. (2008). Overlapping tasks methodology as a tool for investigating language perception (pp. 125–152). In Z. Breznitz (ed.), Brain Research in Language, Springer.

Stroop, J. R. (1935). Studies of interference in serial verbal reactions. Journal of Experimental Psychology, 18, 643–661.

Telford, C. W. (1931). The refractory phase of voluntary and associative responses. Journal of Experimental Psychology, 14, 1–36.

Tombu, M., & Jolicoeur, P. (2003). A central capacity sharing model of dual-task performance. Journal of Experimental Psychology: Human Perception and Performance, 29, 3–18.

Vachon, F. & Jolicoeu, P. (2012). On the automaticity of semantic processing during task switching. Journal of Cognitive Neuroscience, 24, 611–626.

Welford, A. T. (1952). The “psychological refractory period” and the timing of high speed performance – A review and a theory. British Journal of Psychology, 43, 2–19.

West, W.C. & Holcomb, P.J. (2000). Imaginal, semantic, and surface-level processing of concrete and abstract words: an electrophysiological investigation. Journal of Cognitive Neuroscience, 12, 1024–1037.

Ziegler, J. C., & Jacobs, A. M. (1995). Phonological information provides early sources of constraint in the processing of letter strings. Journal of Memory and Language, 34(5), 567–593.

Ziegler, J. C., Tan, L. H., Perry, C., & Montant, M. (2000). Phonology matters: The phonological frequency effect in written Chinese. Psychological Science, 11, 234–238.

